# Roflumilast, a Phosphodiesterase-4 Inhibitor, Ameliorates Sleep Deprivation-Induced Cognitive Dysfunction in C57BL/6J Mice

**DOI:** 10.1101/2021.11.28.470251

**Authors:** Abid Bhat, Muhammed Bishir, Seithikurippu R. Pandi-Perumal, Sulie Chang, Saravana B. Chidambaram

## Abstract

Sleep deprivation (SD) interferes with long-term memory and cognitive functions by over-activation of phosphodiesterase (PDEs) enzymes. PDE4, a non-redundant regulator of the cyclic nucleotides (cAMP), is densely expressed in the hippocampus and is involved in learning and memory processes. In the present study, we investigated the effects of Roflumilast (ROF), a PDE4B inhibitor, on sleep deprivation induced cognitive dysfunction in a mouse model. Memory assessment was performed using a novel object recognition task and the hippocampal cAMP level was estimated by the ELISA method. The alterations in the expressions of PDE4B, amyloid-beta (Aβ), CREB, BDNF, and synaptic proteins (Synapsin I, SAP 97, PSD 95) were assessed to gain insights into the possible mechanisms of action of ROF using the Western blot technique. Results show that ROF reversed SD induced cognitive decline in mice. ROF down-regulated PDE4B and Aβ expressions in the brain. Additionally, ROF improved the cAMP level and the protein expressions of synapsin I, SAP 97, and PSD 95 in the hippocampal region of SD mice. Taken together, these results suggest that ROF can suppress the deleterious effects of SD-induced cognitive dysfunction via the PDE4B-mediated cAMP/CREB/BDNF signaling cascade.

## 1. INTRODUCTION

Sleep plays a regulatory role in maintaining cellular and metabolic homeostasis. Increasing evidence reveals that sleep disturbances affect higher-order brain functions that are linked to various neurological disorders ^1–3^. Sleep deprivation (SD) has a detrimental influence on both social and economic well-being ^4, 5^. Accumulating evidence suggests that SD reduces neurogenesis and protein expression of neurotrophic factors such as CREB and BDNF, which are the crucial regulators for learning and memory and their down-regulation was reported to induce hippocampal atrophy ^6–8^. Functional magnetic resonance imaging (fMRI) and behavioral studies showed that one-night sleep deprivation substantially compromises hippocampal function in humans in turn affecting memory ^9^. Furthermore, sleep disturbance predisposes neuronal accumulation of amyloid-β (Aβ) ^10–12^ in the brain. Positron emission tomography in humans also confirmed that SD and sleep fragmentation are associated with increased deposition of Aβ in the brain ^13, 14^.

A mechanistic research report showed that SD promotes the synthesis and impairs the neuronal clearance of Aβ protein ^15^. Intriguingly, the relationship between SD and Aβ is bidirectional because the other way around increased Aβ deposition impairs slow wave sleep ^16^. Furthermore, Aβ reduces the protein expression of Synapsin I, PSD-95, and SAP-102, which indicates that it eliminates synapses and disrupts neuronal networks ^17–19^. This synapto-toxic effect of Aβ is linked to the reduced expression of NMDA receptors and decreased cAMP content ^20, 21^. The increased accumulation of Aβ and decreased levels of cAMP decreases the release of neurotrophic factors that regulate brain development and synaptic plasticity ^22, 23^.

Phosphodiesterases (PDEs) are a diverse family of enzymes that play a pivotal role in regulating intracellular signaling ^24^. An increased expression of PDE4 enzymes hydrolyse the cAMP into its inactive form 5-AMP, which has been consistently observed in Alzheimer’s disease (AD) patients’ brains ^25^, and subjects with cognitive impairment ^26^ and also in the hippocampal region of SD mice ^27^. PDE4s isoforms such as *PDE4A*, *PDE4B,* and *PDE4D* are expressed in the brain while *PDE4C* is predominantly expressed in testis, and lacks expression in the brain^28^. PDE4B is highly expressed in rodent hippocampal CA2 and CA3 regions, parietal and piriform cortex, and the cerebellar granular layer ^29^. Inhibition of PDE4 improved learning and memory in a mouse model of AD via increasing hippocampal cAMP levels ^25, 30^. Furthermore, inhibition of PDE4 has also restored the deficits in synaptic proteins such as synaptophysin, and PSD 95 ^31, 32^. Specifically, PDE4 inhibition was shown to reverse the cognitive decline induced by muscarinic receptor antagonist ^33^ and also by modulating NMDA receptors mediated transduction mechanisms in rat models ^34^. Albeit, the NMDA does not affect PDE4 expression directly, however, the balance between PDE4 and NMDA mediated adenylyl cyclase plays a pivotal role in the memory process ^34^. Thus, enormous reports suggest that PDE4s are the potential target in neurological disorders drug discovery ^35^.

Roflumilast (ROF), a cAMP-specific PDE4B inhibitor, is approved by USFDA for use in chronic obstructive pulmonary disease (COPD) ^36^. ROF promotes hippocampal neuron viability ^37^ and improves memory in rodents and monkeys at the non-emetic doses ^38^. In a clinical study, it is observed that acute administration (3 days) of ROF improves learning and memory in healthy subjects ^39^. Long-term SD produces an AD-like pathological state, wherein increased neuronal accumulations of Aβ, decreased cAMP, and decreased synaptic protein expressions are well established. This spurs interest to investigate whether PDE4 expression has any correlation with Aβ, CREB, BDNF expression in SD brains, and also to study the effect of PDE4B inhibition, using ROF, on cognitive function in sleep-deprived mice. **Figure.1** explains the flow of the present study’s experimental design.

**Figure 1:**
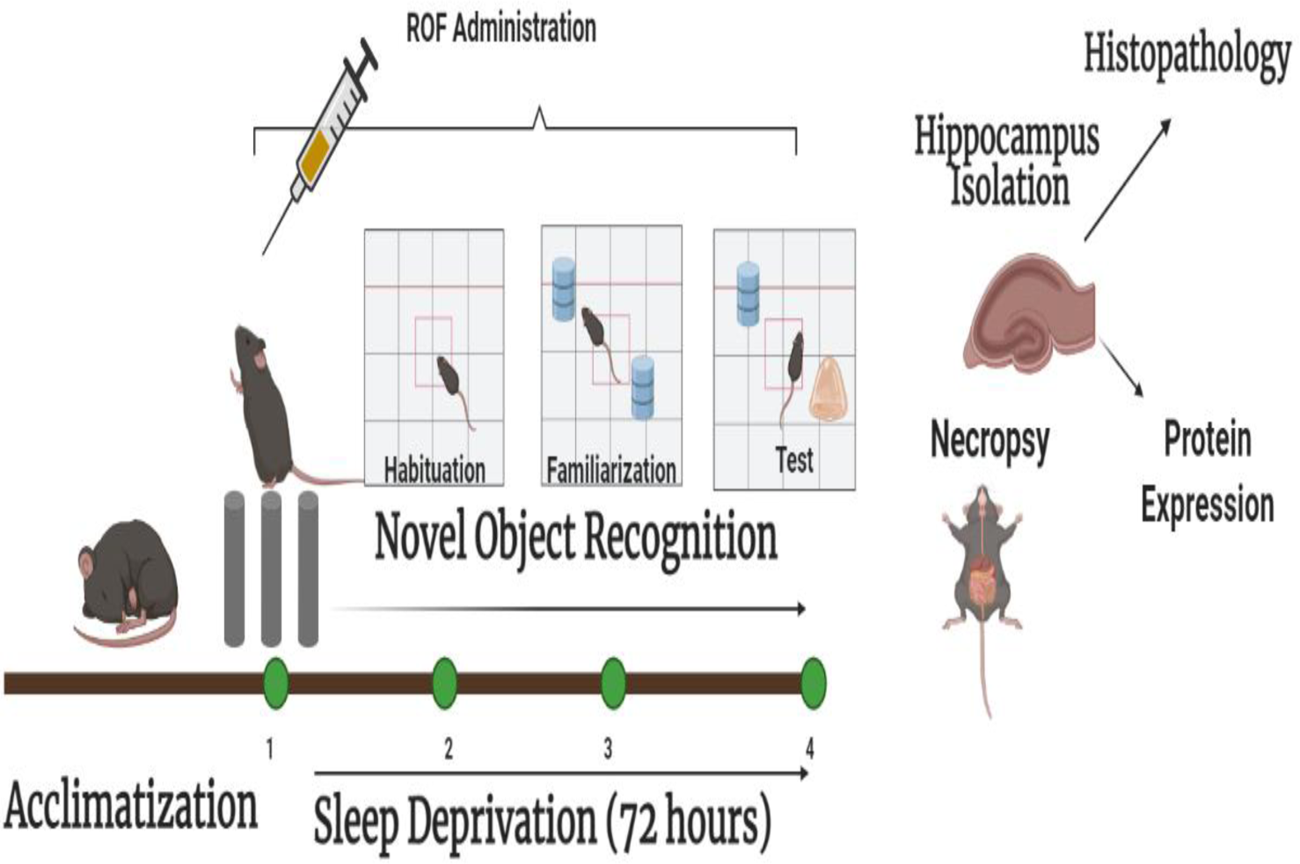
Schematic outline of the experimental design

## 2. RESULTS

### 2.1. SD up-regulated PDE4B expression in the mice hippocampal region

In the present study, we assessed the impact of SD on PDE4B expression in the hippocampal region of mice brains. 72h of continuous SD induced a significant *(p* < 0.05) increase in hippocampal PDE4B expression when compared to the NSD group. The daily dose of ROF down-regulated PDE4B expression when compared to the vehicle-treated SD group. A significant *(p* < 0.01) decrease in PDE4B expression was found at 3 mg/kg dose of ROF (**Fig 2A**).

**Figure 2:**
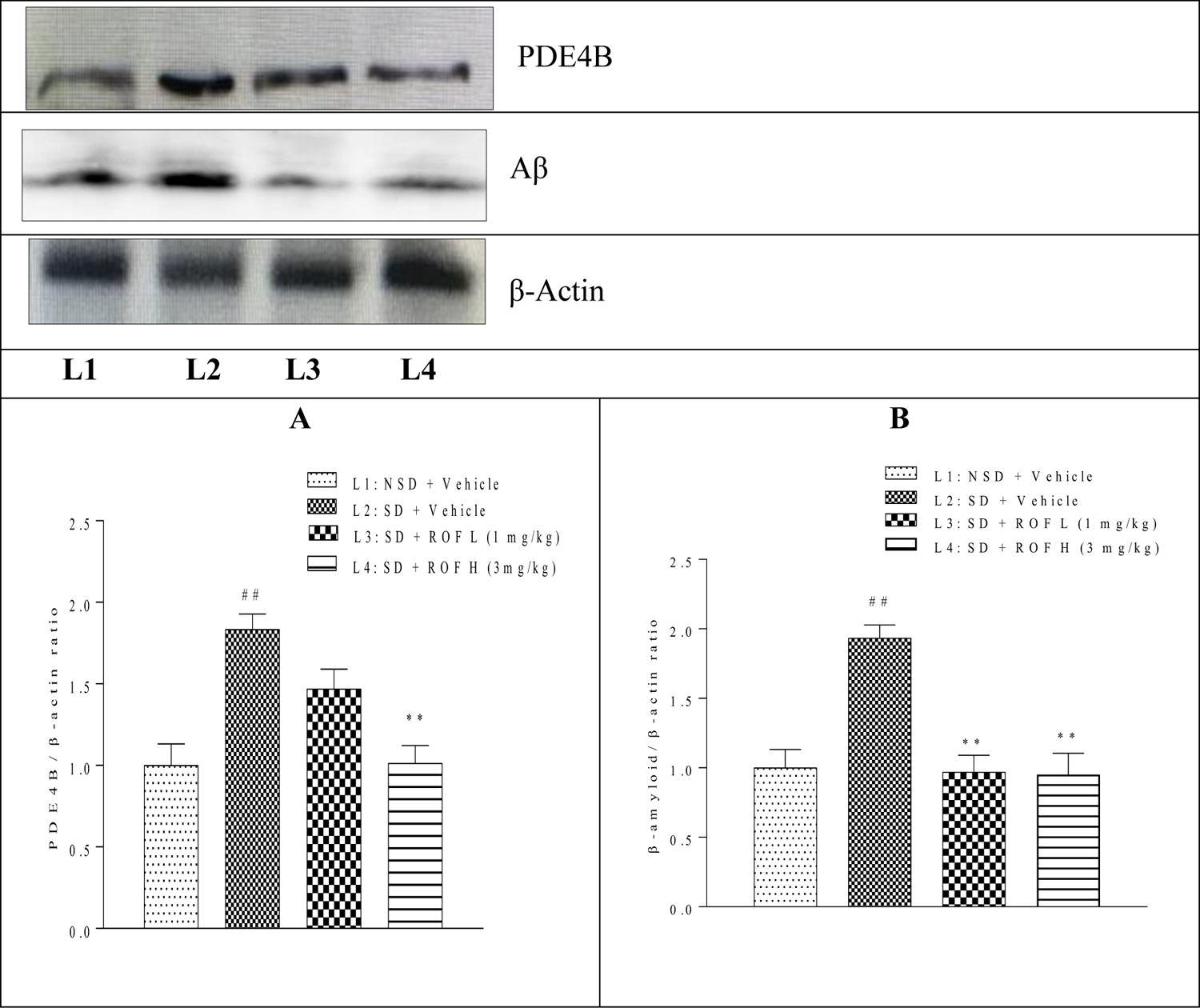
Roflumilast downregulated the protein expression of PDE4B and Aβ in the hippocampus of SD mice. **(A)** Semi-quantification of PDE4B/β-actin expression **(B)** Semi-quantification of Aβ /β-actin expression. Data are presented as the mean ± SEM, ^##^denotes *p*<0.01 versus vehicle-treated NSD group, **denotes *p*<0.01 versus vehicle-treated SD group.

Impaired sleep is linked to AD as sleep plays a role in clearing the metabolic waste from the brain ^11^. We performed a Western blotting assay to study the impact of SD on Aβ expression in mice hippocampal region. Mice subjected to 72h of SD showed a significant (*p*< 0.01) increase in Aβ expression when compared to the non-sleep-deprived mice. Inhibition of PDE4B by ROF significantly (*p*< 0.01) reduced the expression of Aβ in mice when compared with the vehicle-treated SD group.

### 2.2. SD increased Aβ deposition in hippocampal CA1 and DG regions of mice brains and roflumilast decreased Aβ deposition

An increase in Aβ expression was further confirmed by Congo red staining of SD mice brains. We found that vehicle-treated SD mice showed multifocal and intense deposition of Aβ in CA1 and DG regions when compared to the NSD group. ROF-treated SD mice showed very mild deposition of Aβ in CA1 and DG regions when compared to vehicle-treated SD mice brains (**Fig.3**).

**Figure 3:**
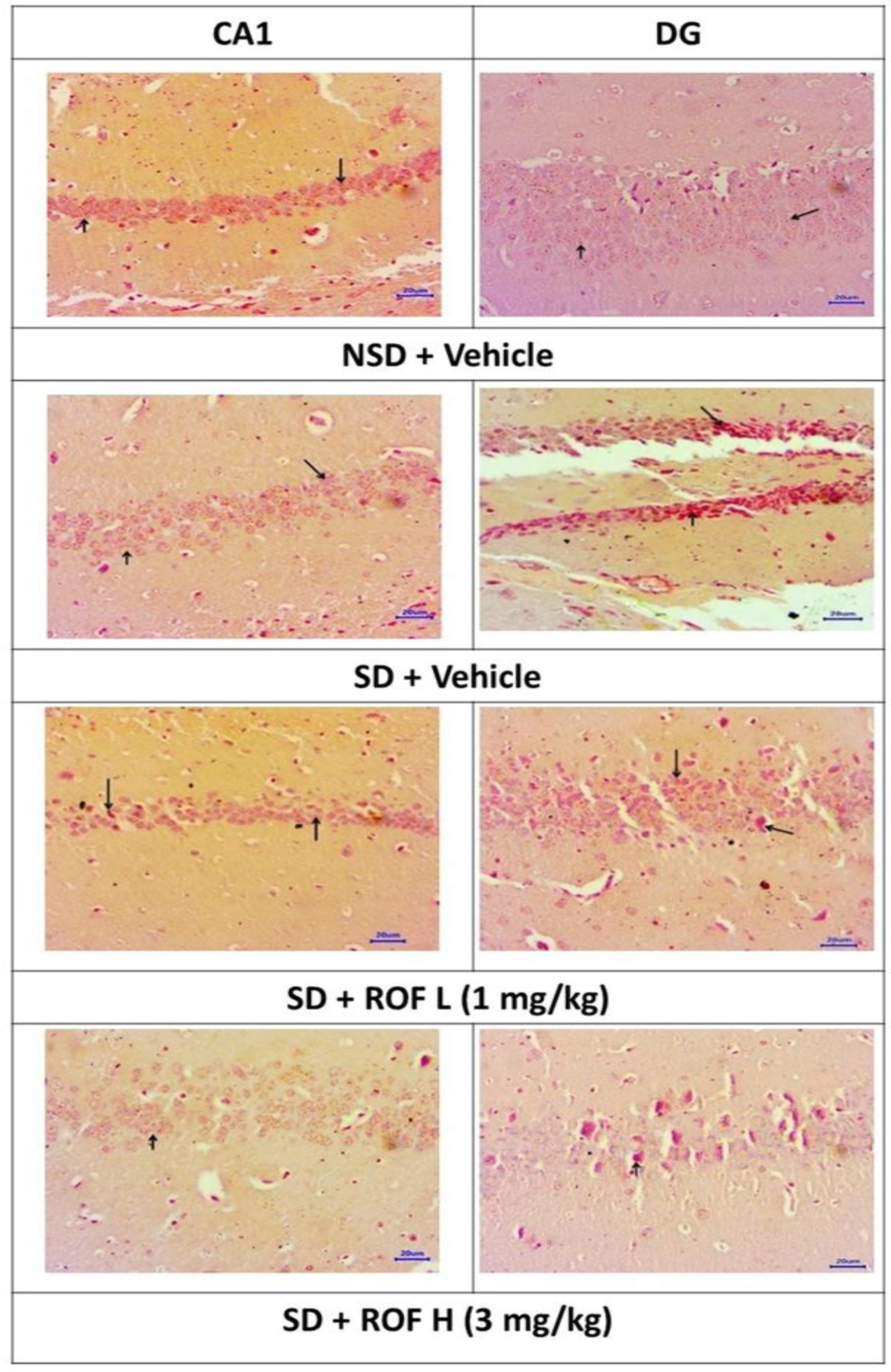
Effect of ROF on Aβ plaques in CA1 and DG regions of the hippocampus in sleep-deprived mice. Arrows denote amyloid-β plaques.

### 2.3. Roflumilast improves hippocampal cAMP levels in sleep-deprived mice

cAMP regulates the learning and memory processes in the brain. The decline of cAMP levels in the hippocampus impairs memory consolidation ^40^. To determine the impact of SD on cAMP levels in the hippocampus region of mice, we performed an ELISA assay and found that SD significantly *(p* < 0.01) reduced cAMP levels in vehicle-treated mice when compared with the NSD group. Administration of ROF in sleep-deprived mice showed a significant *(p* < 0.001) increase in the levels of cAMP when compared with the vehicle-treated SD group. This indicates that ROF can rescue the cAMP levels in SD mice brains. (**Fig. 4**).

**Figure 4:**
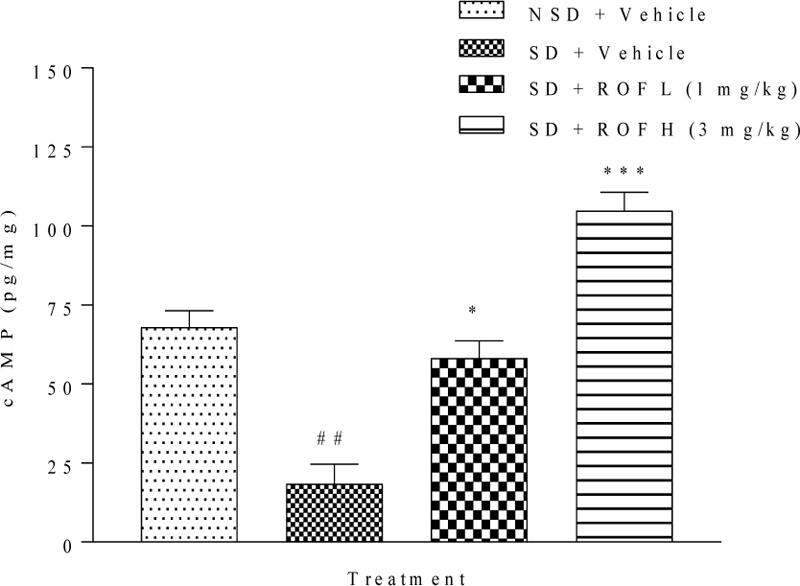
Roflumilast administration restored SD induced decrease in cAMP level in mice. Data are presented as the mean ± SEM. ^##^ denotes *p*<0.01 versus the vehicle-treated NSD group, * and *** denote *p*<0.05 and *p*<0.001, respectively, versus the vehicle-treated SD group.

### 2.4. Roflumilast improved the neurotrophic factors - CREB, and BDNF proteins expression in the hippocampal region of SD mice

Next, we assessed the impact of SD on the protein expression of CREB and BDNF in the hippocampal region of mice brains. We found that 72-h SD produced a significant (*p* < 0.01) decrease in pCREB expression as compared to the NSD group. Administration of ROF restored pCREB expression when compared to the SD control group. A significant (*p* < 0.001) increase in pCREB was observed with ROF treatment when compared with vehicle-treated SD mice (**Fig. 5A**).

**Figure 5:**
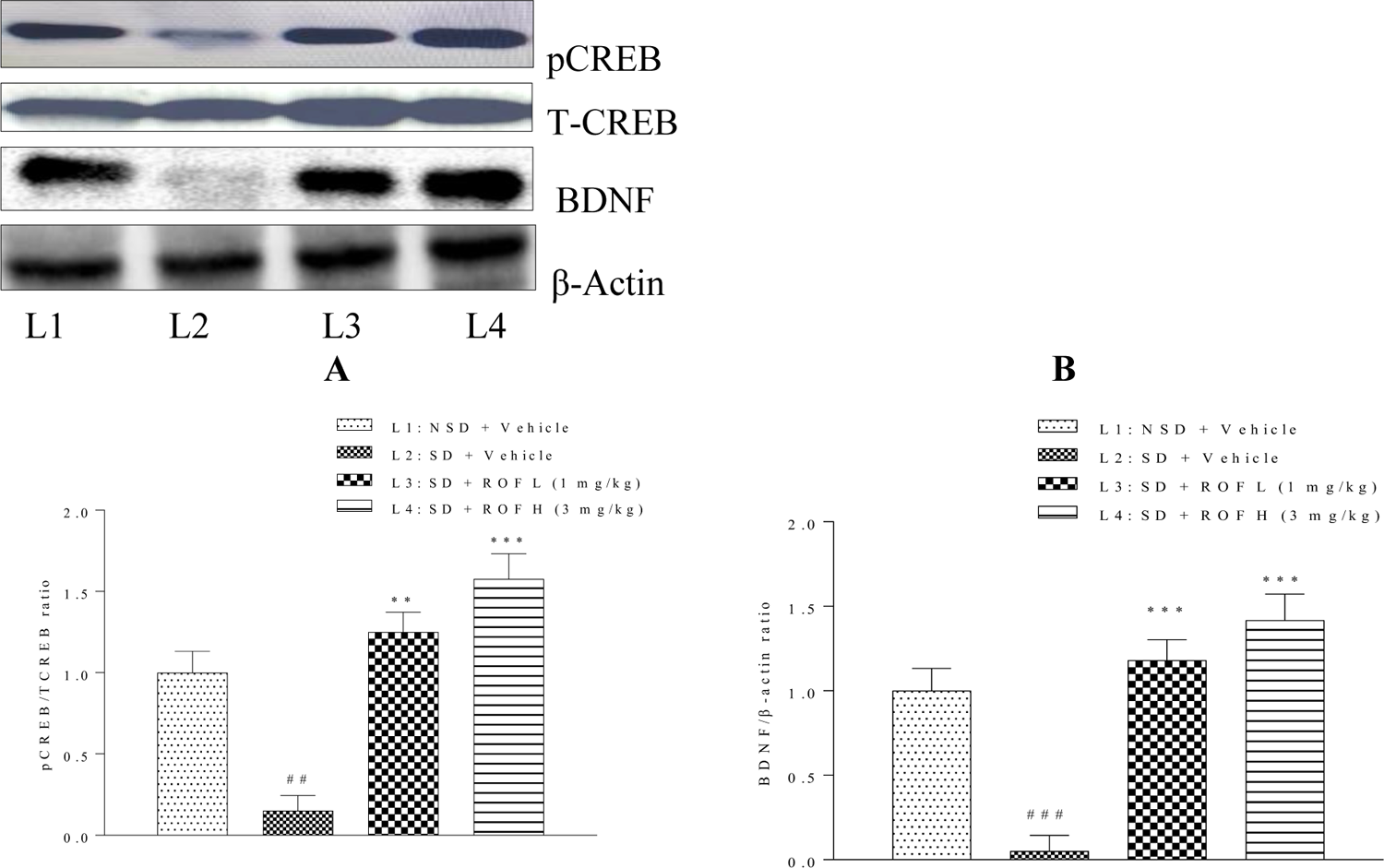
Roflumilast administration increased neurotrophic factors, CREB and BDNF, in sleep-deprived mice brain. **(A) Semi-**quantification of pCREB/TCREB expression **(B)** Semi-quantification of BDNF /β-actin expression. Data are presented as the mean ± SEM, ^##^ denotes p < 0.01, ^###^ denotes p < 0.001 versus vehicle-treated NSD group, ** and *** denotes *p*<0.01, and *p*<0.001, respectively, versus vehicle-treated SD group.

CREB positively influences the expression of BDNF, which is essential in memory consolidation and synaptic functions ^6^. Hence we performed Western blot analysis to detect BDNF expression in the hippocampus region of SD mice. As shown in **Fig. 5B**, SD significantly (*p* < 0.001) decreased the BDNF protein expression when compared with NSD mice. ROF administration in sleep-deprived mice significantly (*p*< 0.001) increased BDNF expression when compared to vehicle-treated SD mice. These data suggest that ROF administration improves the protein expression of neurotropic factors CREB and BDNF in SD mice.

### 2.5. Roflumilast up-regulates the expression of the synaptic proteins in sleep-deprived mice brains

Next, we investigated whether improvement in neurotrophic factors expression has any influence on the expression of the synaptic proteins in the hippocampal region of SD mice. Synapsin I expression decreased significantly (*p*< 0.01) following SD when compared to the NSD control group. ROF administration significantly (*p*< 0.01) up-regulated the expression of Synapsin I when compared to vehicle-treated SD mice (**Fig 6**). SAP 97 regulates synaptic plasticity by controlling the distribution of NMDA- and AMPA-type glutamate receptors ^41^. We found a significant decrease in the expression of SAP-97 in SD mice which was reversed (p<0.001) with ROF treatment (**Fig 6**).

**Figure 6:**
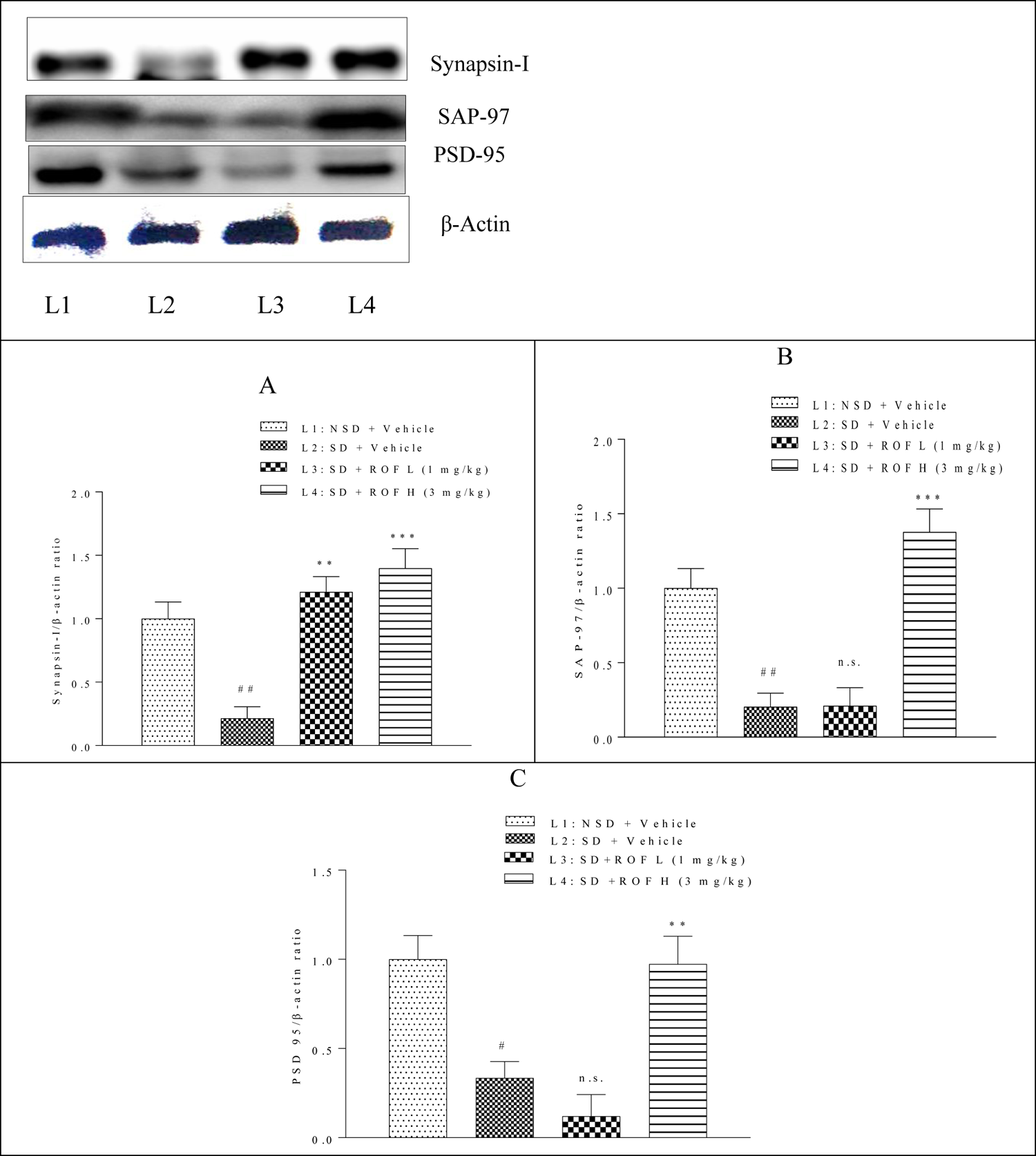
Roflumilast upregulated the expression of synaptic proteins in SD mice brain. **(A)** Semi-quantification of Synapsin-I/β-actin expression. **(B)** Semi-quantification of SAP-97/β-actin expression. **(C)** Semi=quantification of PSD-95/β-actin expression. Data are presented as the mean ± SEM. ^#^ denotes *p*<0.05, ^##^ denotes *p*<0.01 versus vehicle-treated NSD group, ** and *** denotes *p*<0.01 and *p*<0.001, respectively, versus vehicle-treated SD group, n.s. denotes non-significant.

Concurrently, we investigated the impact of SD on the post-synaptic density protein, PSD-95, expression. 72 h of SD caused a significant (*p*< 0.01) decrease in PSD-95 expression when compared to NSD mice. Administration of ROF significantly up-regulated (*p*< 0.01) PSD95 expression in SD mice (**Fig 6**). These results imply that PDE4B inhibition can improve synaptic functions, which could be corroborated by the restoration of neurotrophic factors, at least partly, in SD.

### 2.6. Roflumilast restores sleep-deprivation induced cognitive dysfunction in mice

Recognition memory in mice was assessed using NORT. Sleep-deprived mice showed a significant (*p* < 0.05) decrease in time spent with the novel object when compared to NSD mice. Administration of ROF significantly (1mg/kg; *p* < 0.05, 3mg/kg; *p* < 0.01) increased time spent with the novel object by SD mice (**Fig. 7a**). SD significantly (*p* < 0.01) decreased the discrimination index, but ROF significantly (1mg/kg; *p* < 0.05, 3mg/kg; *p* < 0.01) increased the discrimination index in SD mice (**Fig. 7b**). Decreased Aβ, and increased CREB / BDNF and synaptic proteins (synapsin-1, SAP-97, and PSD95) expression are found to corroborate to the improved memory in SD mice.

**Figure 7:**
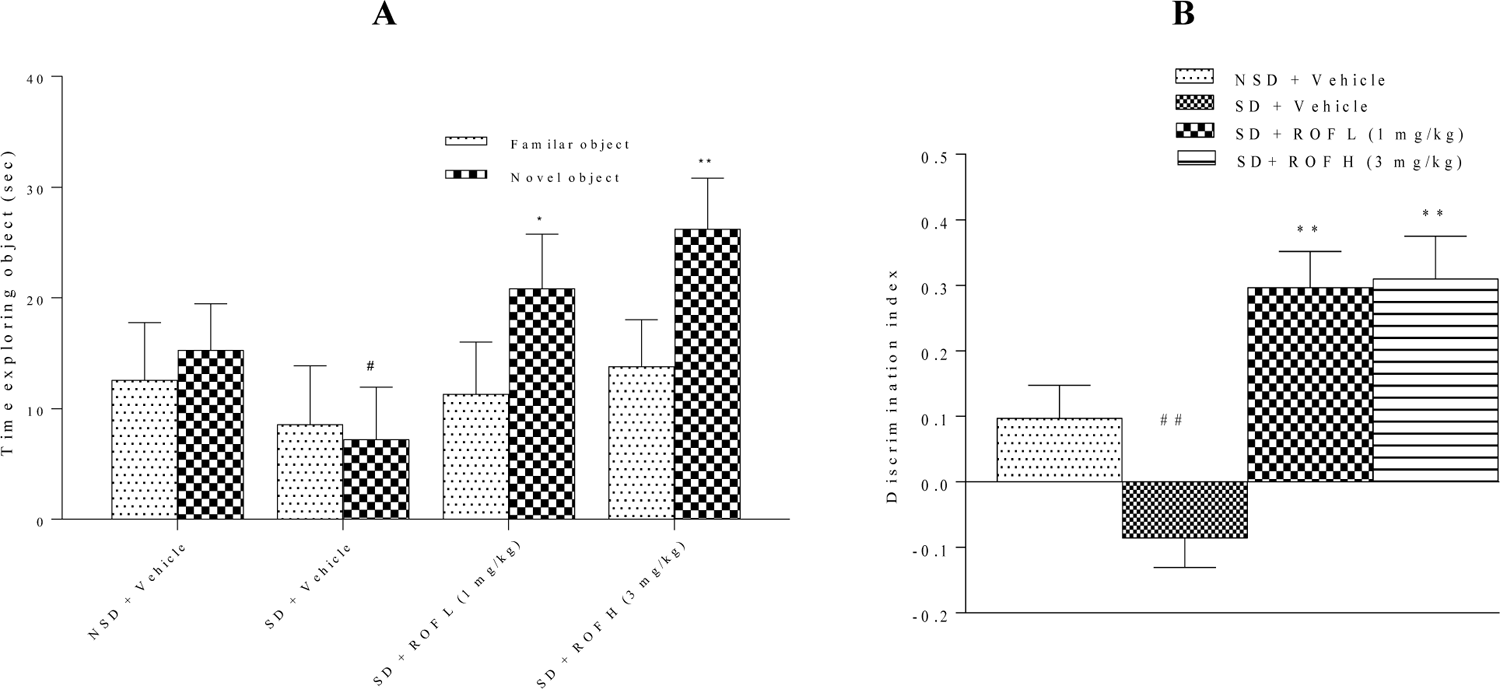
Roflumilast improved recognition memory in NORT in sleep-deprived mice. (A) time exploring the familiar and novel object (B) Discrimination index. Data are presented as the mean ± SEM. ^#^ and ^##^ denotes *p* <0.05, and *p*<0.01 respectively, versus the NSD group and * and *** denote *p* <0.05, and *p*<0.01, respectively, versus the SD group

### 2.7. Roflumilast prevented the morphological changes of hippocampal neurons in sleep-deprived mice

SD induced multifocal moderate neuronal degeneration in the CA1 and DG regions of the hippocampus. As shown in **Fig 8**, neurons in CA1 and DG regions in vehicle-treated SD mice showed a decrease in purple-stained Nissl granules and increased pyknotic nuclei in the perikarya when compared to NSD mice. ROF-treated mice showed reduced morphological changes and had regularly shaped neuronal bodies in CA1 and DG regions when compared to SD mice (**Fig 8**).

**Figure 8:**
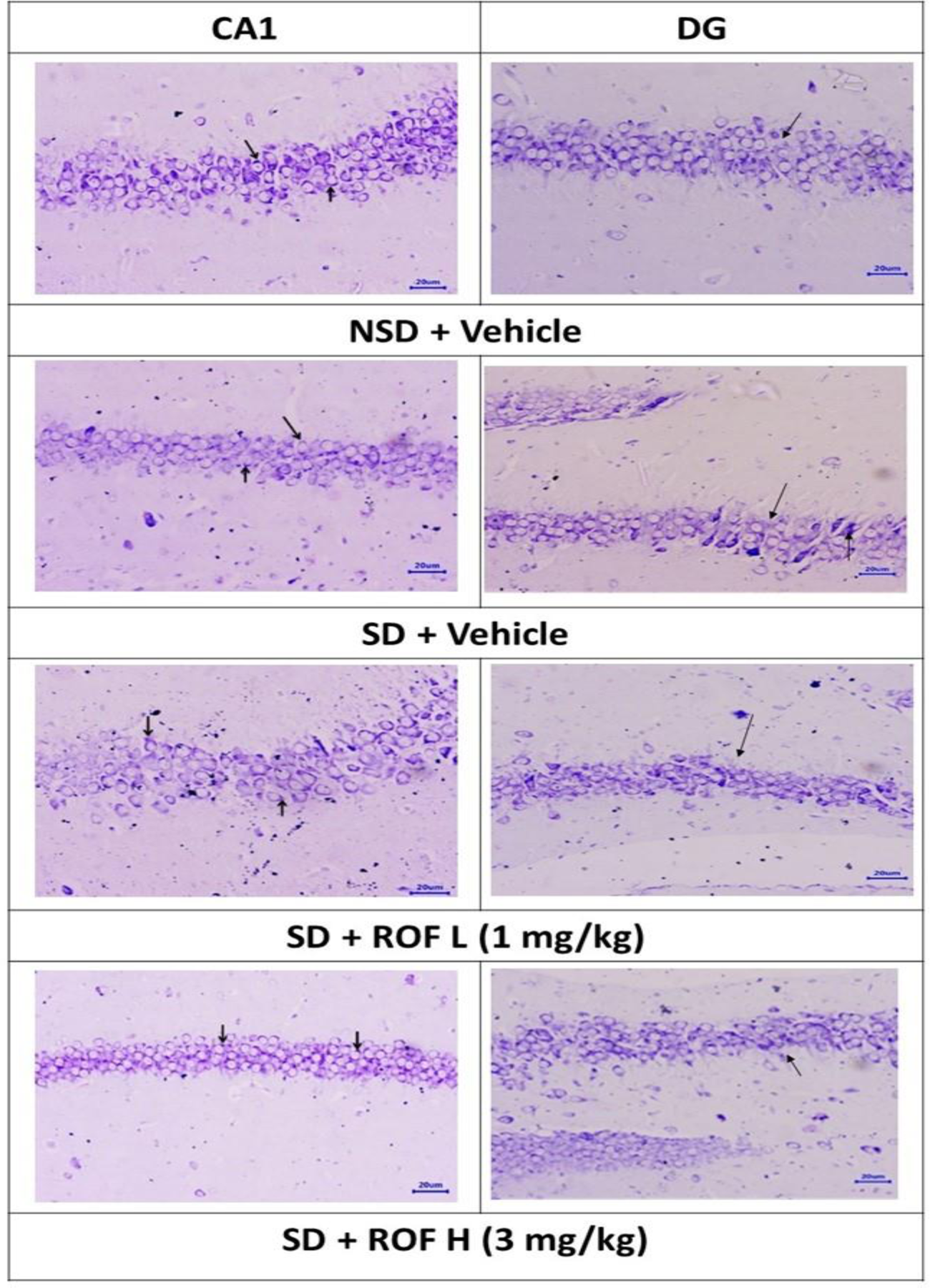
Cresyl violet staining of CA1 and DG regions of the hippocampus of SD mice. Arrows denote pyknotic nuclei.

## 3. DISCUSSION

The present study reveals that alleviation of Aβ pathology, cAMP signaling, and synaptic proteins expression by ROF via PDE4B inhibition is the key mechanism in cognitive restoration in SD mice.

SD aggravates Aβ plaque levels in AD transgenic mouse model ^42^. A recent study has shown that chronic sleep restriction (3 h per day, 5 days per week, for 4 weeks) increases the hippocampal accumulation of Aβ plaques in mice brains, which was corroborated by the cognitive decline ^43^. Aβ deposition in cortical and hippocampal regions initiates inflammatory responses, synaptic dysfunctions ^44^, and neuronal apoptosis ^45^. Aβ also compromises the cAMP-response element-binding protein (CREBP) signaling in neurons suggesting that multiple factors contribute to neuronal damage in SD ^46^.

On the other hand, increased expression of PDE4D is reported to associate with Aβ plaque pathology and memory loss ^47^. Studies using PDE4D knockout mice showed improved memory and reduced neuroinflammation and β-amyloidosis ^48–50^. In the present study, we found that 72h SD increased the protein expression and/or neuronal deposition of Aβ in hippocampal neurons of mice. Earlier, in vivo studies and clinical trials have shown that alleviating cAMP signaling through PDE4 inhibition improves cognitive functions ^51, 52^ and reduced Aβ expression in mice ^53^. The present study reports an increased PDE4B and Aβ expression in SD mice brains with a significant decrease in cAMP levels. Treatment with ROF reversed the molecular changes including Aβ pathology and improved cognitive performance in SD mice which could be related to PDE4B inhibition.

Synapse dysfunction is an immediate consequence of increased Aβ deposition ^54^. PSD-95 is an essential regulator of synaptic strength and plasticity. SAP-97 facilitates synaptic vesicles biogenesis, and neurotransmitters release ^55^. Synapsin I regulates neurotransmitters release and its down-regulation is shown to impair neurogenesis and synaptic plasticity ^56, 57^. SD causes synaptic damage in the hippocampal region by reducing the expression of presynaptic and postsynaptic proteins ^58–60^. Recently, we have shown that ROF improves synaptic protein expression using human neurons (in vitro model) exposed to the neurotoxin quinolinic acid ^31^.

In the present in vivo study, we report that SD down-regulates synaptic proteins (SAP-97, Synapsin-I, and PSD-95) in the hippocampal region of mice. Nevertheless, administration of ROF increased SAP-97, Synapsin-I, and PSD-95 expression, which indicates that PDE4B plays a role in synaptic protein expression.

Inhibition of cAMP signaling in the hippocampal region is reported to impair the consolidation of long-term memory in mice ^61^. SD reduces the phosphorylation of CREB in the hippocampus and affects the protein expression of the BDNF ^6^. In the present study, ROF administration restores the cAMP/CREB/BDNF signaling cascade in SD mice. These results are in agreement with earlier studies that report that inhibition of PDE4 improves memory consolidation in rodent models via the cAMP/CREB/BDNF cascade ^33, 62^.

Histopathological examination using Congo red staining showed intense deposition of Aβ plaques in the CA1 and DG regions of the hippocampus in SD mice. Further, cresyl violet staining showed neuronal death in the hippocampus region of SD mice. Recent studies have shown that SD increases cytokine production, and microglial activation and initiates neuronal apoptosis, causing lesions in the hippocampus of mice ^63, 64^. Increased PDE4B expression is reported to trigger inflammation and microglial reactivity ^65^. Treatment with ROF reduced Aβ plaques and also increased the Nissl granules in the hippocampal regions in SD mice. Inhibition of PDE4 transiently increases cAMP and stimulates cAMP signal transduction, which in turn, reduces the levels of pro-inflammatory and early apoptosis factors ^66^. This could be a possible mechanism for the observed neuroprotection with ROF in SD

## 4. CONCLUSION

From the current investigation, we provide molecular, behavioral, and histopathological evidence that substantiates the protective effects of Roflumilast against SD-induced cognitive dysfunction in a mouse model. Roflumilast administration improves recognition memory via PDE4B inhibition which was found to be mediated via cAMP/CREB/BDNF signaling and downregulation of Aβ pathology in SD mice. Additionally, further studies to understand the effect of Roflumilast on NMDA activity and autophagy in chronic sleep restriction are under investigation in our lab.

## 5. MATERIALS AND METHODS

### 5.1. Animals

Male C57BL/6J mice (25-30 g) were obtained from M/s. Adita Biosys Private Limited, Tumakuru, Karnataka, and housed in CPT, JSS Academy of Higher Education & Research, Mysuru, Karnataka. Animals were housed in groups (5 mice/cage) in polypropylene cages under an ambient temperature of 19-25^°^C and 40-65% relative humidity, with a 12-h light/dark artificial light cycle. Animals were provided with standard rodent feed and purified water ad libitum. Animals were acclimatized for 7 days to the laboratory conditions prior to initiation of the experiments. Animal experiments were performed in full compliance with the guidelines of the “Guide for the Care and Use of Laboratory Animals” National Research Council, 2011). Institutional Animal Ethics Committee (IAEC), Central Animal Facility, JSS AHER, Mysuru, India, has approved the study protocol (JSSAHER/CPT/IAEC/014/2020).

### 5.2. Reagents and antibodies

Roflumilast, Cresyl violet, and Congo red stains were purchased from Sigma Aldrich (India). cAMP ELISA kit was purchased from Cayman (Ann Arbor, MI, USA). Anti-PSD95 (sc-32290), Anti-SAP97 (sc-9961), Anti-Synapsin-I (sc-376623), Anti-BDNF (sc-65514), Anti-CREB (sc-377154), Anti β-Amyloid (sc-28365) were procured from Santa Cruz Biotechnology, CA, USA. Anti-PDE4B (NB100-2562) was purchased from Novus Biologicals, United States. All other reagents and chemicals were analytical grade.

### 5.3. Roflumilast treatment

Roflumilast [ROF (L): 1 mg/kg and ROF (H): 3 mg/kg] was freshly prepared in 0.5% carboxy methyl cellulose (CMC) and administered intraperitoneally once a day for three days. The doses of ROF was selected based on earlier published report^67^.

### 5.4. Sleep deprivation method

Sleep deprivation of mice was done via a modified multiple platform method^68^. Briefly, a mouse was placed on a cylindrical platform (8.5 cm in height and 2.5 cm in diameter). Eight platforms were kept in a cage approximately 10 cm apart inside a cage filled with water up to 2 cm from the bottom. Water was changed twice a day. The size of the platforms permits the mouse to sit but not lie down. Furthermore, mice could easily move between the platform but could not stretch across the platforms. This method is reported to eliminate REM sleep ^69^. Non-sleep-deprived (NSD) animals were kept in normal polypropylene cages in the same room. During sleep deprivation, mice had access to food and drinking water. Animals were sleep-deprived for 72 hours.

### 5.5. Groups and Treatments

Animals were randomized in to four groups based on the stratified bodyweight viz Group 1-: Non-sleep-deprived (NSD) mice which received 0.5% carboxy methyl cellulose [0.5% CMC; n=10]; Group – II: SD which received 0.5% CMC; n=10; Group III and IV: SD which received ROF (L) (1 mg/kg b.wt; n=10) and ROF (H) (3 mg/kg b.wt; n=10), respectively.

### 5.6. Novel object recognition test

A novel object recognition test was performed to access the recognition memory in rodents ^70^. Briefly, the mouse was individually habituated to exploring the empty apparatus for 10 min (2^nd^ day of treatment). During the acquisition trial (T1), two similar objects were placed inside the apparatus and the mouse was allowed to explore the objects for 3 min. After the acquisition trial, the mouse was transferred to its home cage. The discrimination trial (T2) was done twenty-four hours later (4th day). Two different objects, a familiar object, and a novel object were placed in the exploration area. The time spent by the animal exploring the two objects during T2 was recorded and the discrimination index (DI) was calculated, as per the following formula. DI=RI/ (Time spent in exploring novel object + Time spent in exploring familiar object). Recognition Index = Time spent in exploring novel objects-Time spent in exploring the familiar objects. This test was repeated for all the animals in all the cages one at a time. All behavioral assessments were done between 10.00 am to 1.00 pm.

### 5.7. Measurement of cAMP content

The hippocampal region of the mouse was isolated and homogenized using 0.1M HCl (10% tissue homogenate). Homogenates were centrifuged for 10 min at 1500× g at 4°C, and the supernatants were stored at 4°C. cAMP levels were determined by a cAMP enzyme immunoassay kit following the manufacturer’s instructions (Cayman Chemical Co., Ann Arbor, MI, USA)

### 5.8. Western blot

Following the behavioral assessment, the animals were euthanized to collect the brain and stored at −80°C. Hippocampal region was isolated, and homogenates were prepared with radioimmunoprecipitation assay buffer (RIPA) buffer (50 mM Tris, pH 7.4, 150 mM NaCl, 1% NP-40, 5 mM EDTA, 0.5% sodium deoxycholate, 0.1% SDS, 50 nM sodium fluoride, 1 mM sodium vanadate) containing a cocktail of protease inhibitor (Sigma Aldrich, MO, USA). Total protein concentrations of the samples were determined by the Pierce™ bicinchoninic acid (BCA) protein assay (Thermofisher Scientific, USA), and samples were aliquoted and stored at −80°C until further use. Sample proteins (20 μg) were separated by using 10% bis-tris-SDS-PAGE (electrophoresis). Resolved proteins in the gels were transferred onto polyvinylidene difluoride (PVDF) membranes (Biorad, USA) and electroblotted. Membranes were blocked overnight with 5% non-fat skimmed milk in Tris-Buffered Saline and Tween 20 (TBST) at 4°C. This was followed by a 4-h incubation with the primary antibodies (PDE4B (1:1000), CREB (1:1000), BDNF (1:1000), β-Amyloid (1:1000), PSD-95 (1:1000), Synapsin-I (1:1000), SAP 97 (1:1000) at room temperature. The membranes were rinsed with TBST (3 washings for 10 min each), followed by incubation with the secondary antibodies (HRP conjugated anti-mouse or anti-rabbit IgG) for 1h at room temperature and washed with TBST (3 washings for 10 minutes each). Bands were detected using SuperSignal West Pico PLUS Chemiluminescent Substrate (Thermo Scientific, USA). Densitometric measurement of bands was done using ImageJ (NIH software). For Western blot analysis, the signal intensity (integrated density value, IDV) of PDE4B, BDNF, β-Amyloid, PSD-95, Synapsin-I, and SAP 97 was normalized against the IDV of internal control β-actin, while pCREB was normalized with total CREB and histogram was plotted.

### 5.9. Histopathology

Whole-brain was stored in buffered 10% formalin for 48h. Coronal sections (3-5 µm) of the hippocampus region were cut using a microtome. The hippocampal region of the brain was used for histopathological analysis. Sections were mounted on a slide, washed, and dehydrated with 95% ethanol.

### 5.10. Cresyl violet staining

Coronal sections (3-5 µm) of the hippocampus region were washed with xylene and followed by five times washing with water at 5 min intervals. The samples were stained using 0.2% cresyl violet dye for 30 min. The prepared slides were examined under the microscope by a pathologist, blinded to the treatment protocol, for histopathological examination.

### 5.11. Congo red staining

Congo red staining was performed to detect the deposition of amyloidβ plaques in mice hippocampus. Coronal sections (3-5 µm) of the hippocampus region were stained with 1% Congo Red stain for 30 min. Amyloidβ plaques were observed under a microscope.

### 5.12. Statistical analysis

Data are presented as mean ±SEM. The difference between time spent exploring the novel object versus a familiar object during the discrimination trial was calculated for each group and the level of significance was analyzed using a two-sided student’s t-test. For other parameters, group means differences were analyzed using a one-way ANOVA test followed by Tukey’s multiple comparison test as post hoc. Graphs were plotted using GraphPad Prism version 7.04 with a *p-value* ≤ 0.05 is considered significant.

## Supporting information

supplementarr antibody

## ACKNOWLEDGMENTS

AB wishes to thank the Indian Council of Medical Research (ICMR, New Delhi, India) for awarding him a Senior Research Fellowship (45/07/2019/PHA/BMS) during his Ph.D. degree. The authors would like to thank Shri Panchaksharappa Gowda DH, Biostatistician, JSS College of Pharamcy, Mysuru, for his assistance with the biostatistical analysis. All of the authors thank their respective institutions for the administrative and technical assistance provided during the study duration and manuscript preparation.

## AVAILABILITY OF DATA AND MATERIAL

The data sets generated and/or analyzed during the study are available in CPT, JSSAHER.

## ETHICS APPROVAL

The animal experiment protocol was approved (JSSAHER/CPT/IAEC/014/2020) by the Institutional Animal Ethics Committee, JSS AHER, Mysuru, India.

## AUTHOR CONTRIBUTIONS STATEMENT

**AB:** Methodology, Investigation, Data curation, Formal analysis, Writing – original draft, **MB**: Methodology, Investigation, **SRP:** Writing – review & editing, **SC:** Writing – review & editing, **SBC**: Conceptualization, Project administration, – review & editing. The manuscript has been read and approved by all authors.

