## supplementarr antibody for "Roflumilast, a Phosphodiesterase-4 Inhibitor, Ameliorates Sleep Deprivation-Induced Cognitive Dysfunction in C57BL/6J Mice"

**List of antibodies used**

| **Antibody** | **Company** | **Catalog** | **Dilution** |
| --- | --- | --- | --- |
| PDE4B | Novus Biologicals | NB100-2562 | 1:1000 |
| CREB | Santa Cruz Biotechnology | sc-377154 | 1:1000 |
| BDNF | Santa Cruz Biotechnology | sc-65514 | 1:1000 |
| β-Amyloid | Santa Cruz Biotechnology | sc-28365 | 1:1000 |
| PSD-95 | Santa Cruz Biotechnology | sc-32290 | 1:1000 |
| Synapsin-I | Santa Cruz Biotechnology | sc-376623 | 1:1000 |
| SAP 97 | Santa Cruz Biotechnology | sc-9961 | 1:1000 |
